# Combining circulating tumor cell and circulating cell free DNA analyses enhances liquid biopsy sensitivity in detecting high grade serous tubo-ovarian carcinoma

**DOI:** 10.1101/2025.10.13.681971

**Authors:** Beatrice Cavina, Simona Corrà, Camelia Alexandra Coadă, Monica De Luise, Silvia Lemma, Sara Coluccelli, Antonio De Leo, Stella Di Costanzo, Francesco Mezzapesa, Giulia Girolimetti, Pierandrea De Iaco, Anna Maria Porcelli, Anna Myriam Perrone, Dario de Biase, Giuseppe Gasparre, Ivana Kurelac

## Abstract

**Background:** Liquid biopsy is a promising strategy for detecting and monitoring neoplastic diseases, with circulating tumor cells (CTCs) and circulating tumor DNA (ctDNA) being the most common objects of investigation. Most analyses have focused on these biomarkers separately, and simultaneous detection has not yet been attempted in high grade serous tubo-ovarian carcinoma (HGSOC). The aim of our study was to assess whether the tandem CTC/ctDNA approach increases HGSOC detection efficiency of peripheral blood liquid biopsy.

**Methods:** For CTC detection, by using healthy donor samples spiked with known cancer cell numbers, we tested gene expression assays and *TP53* next-generation sequencing (NGS). The latter was also applied for ctDNA detection where analytical validity was ensured by calculating the optimal variant allele frequency (VAF) threshold for mutation calling. The clinical validity of the assays was then verified on two HGSOC cohorts and respective controls. Standard 2x2 contingency tables and Wilson method were used to evaluate clinical validity, by calculating specificity, sensitivity, and accuracy with 95% confidence intervals (CI).

**Results:** High analytical sensitivity and specificity were found for both gene expression and *TP53* NGS based CTC detection, as these assays specifically detected as few as five cancer cells spiked in control sample. Regarding clinical validity, the gene expression-based CTC detection showed 0.48 accuracy, 13.3% sensitivity, and 100% specificity, whereas *TP53* sequencing demonstrated better assay performance (0.77 accuracy, 46.7% sensitivity, 100% specificity). For circulating cell-free DNA (cfDNA) analysis, we first identified 0.31% VAF cut-off for accurate ctDNA *TP53* mutation calling. Subsequent clinical validity assessment showed solid performance efficiency of the ctDNA based liquid biopsy (0.71 accuracy, 60% sensitivity, and 100% specificity), outperforming the CTC detection methods.

Importantly, the tandem ctDNA/CTC analysis improved disease detection rate in both HGSOC cohorts, allowing to achieve, respectively, 73.3% and 93.3% sensitivity. Interestingly, *TP53* NGS revealed CTC private variants, and shared ctDNA/CTC mutations undetected in the primary tissue, highlighting the ability of the dual-analyte approach to capture tumor heterogeneity and allow mutation cross-validation.

**Conclusions:** Our study reveals the complementary value of simultaneous CTC and cfDNA analysis in HGSOC, advancing the translational potential of liquid biopsy integration for the management of this disease.

## Background

(PB) Liquid biopsy, in terms of searching for tumor derived material in peripheral blood samples, has emerged as a revolutionary tool in oncology. It is a minimally invasive approach which allows easily repeatable sampling to monitor disease progression and holds great potential to be implemented for cancer screening [1]. Introducing efficient liquid biopsy in ovarian cancer (OC) setting would be particularly groundbreaking, especially in high grade serous tubo-ovarian carcinoma (HGSOC), the most common and most aggressive histotype, which in 80% of cases is diagnosed at advanced stages due to unspecific and late presentation of symptoms [2,3]. The HGSOC diagnosis relies on histological evaluation of invasive tumor biopsy and, currently, no existing general population screening protocol is available. Finding an appropriate method for recognizing the disease at earlier stages is the most cogent issue in the field, as catching HGSOC at stage I guarantees 91.7% 5-year survival rate, compared to 31.8% when it is diagnosed at stages III-IV [2]. Liquid biopsy approaches offer a promising solution for this challenge, as well as for improvement of patient follow-up in terms of relapse and therapy-response prediction, while reducing the use of invasive procedures. However, liquid biopsy workflows in HGSOC are still at an explorative phase.

(CTCs) Among others, circulating tumor cells and circulating tumor DNA (ctDNA) have been explored in the context of OC, mainly regarding progression monitoring. In particular, CTCs have been associated with worse overall and progression-free survival of OC patients [4,5], but lack of standardized methods for their detection has, so far, prevented such analysis from being implemented into everyday practice. Apart from the general difficulty in discriminating scant tumor-derived material in blood specimens, detection of CTCs in OC is additionally challenged by the high molecular heterogeneity of this disease [6]. For example, whereas positivity for epithelial cell adhesion molecule (EpCAM) is an efficient marker to pinpoint CTCs in breast or lung malignancies [7,8], in OC patients, EpCAM+ cells circulating in PB are not necessarily tumor derived [8]. One of the strategies designed to circumvent the issue of heterogeneity envisions molecular marker agnostic CTC enrichment, such as Parsortix, and subsequent multiple gene expression analysis [9,10]. However, there is no consensus on which gene panel secures the highest specificity and sensitivity, and the efficiency of different enrichment methods is rarely evaluated in a single study [6,11].

(cfDNA) Regarding circulating cell-free DNA analysis, works on HGSOC exploited *TP53* mutations as a distinctive marker of ctDNA [12–15], since this gene is mutated in nearly all cases of HGOSC [16,17]. However, hotspot analysis is not feasible for tumor suppressors unless the variant is known *a priori*, and untargeted next-generation sequencing (NGS) of cfDNA often detects variant allele frequencies (VAF) <0.5, which are not considered to be highly specific markers [14]. Indeed, there is no generally accepted VAF cut-off value for ctDNA *TP53* variant calling specifically in the context of HGSOC.

Interestingly, oncologic liquid biopsy studies have up to date been mainly focusing on either CTC or ctDNA detection separately, rarely performing multianalyte assessment in the same blood sample, even though it is emerging that the two analytes are not redundant biomarkers, but provide complementary information [18,19]. In this scenario, the clinical impact of tandem CTC and ctDNA detection has never been explored in HGSOC and still lacks a formal evaluation.

To converge towards the development of efficient liquid biopsy test for HGSOC detection, the aim of this study was to prove that including both analytes in the workflow increases sensitivity. We set up nucleic acid-based assays for CTC and ctDNA identification and evaluated their analytical and clinical validity, which are critical parameters in any liquid biopsy assay development [1]. By applying these optimized assays on two HGSOC patient cohorts, we demonstrate for the first time that, compared to single analyte approaches, the combination of CTC and ctDNA analyses improves the disease detection rate.

## Methods

### Aim, design and setting of the study

(Figure 1A) The primary aim was to demonstrate that CTC analysis coupled to ctDNA assay increases liquid biopsy-based disease detection rate in HGSOC patients compared to single analyte approach. The secondary aim was to improve standardization of liquid biopsy for HGSOC. We first developed specific assays for each analyte of interest, ensured the analytical validity and then evaluated the clinical validity. For CTC analysis, we compared two enrichment methods (Parsortix and CD45 negative selection) and tested three nucleic acid-based detection techniques: (i) gene expression assays based on cDNA preamplification coupled to quantitative Real Time-PCR (qRT-PCR); (ii) gene expression detection via digital PCR (dPCR); and (iii) CTC mutation identification by *TP53* NGS with Ion AmpliSeq HD laboratory-developed panel (Thermo Fisher Scientific). Analytical validity of each CTC detection approach was evaluated by using healthy donor PB spiked with known numbers of cancer cells, whereas clinical validity was evaluated by analyzing HGSOC patients and healthy controls. For ctDNA detection, *TP53* NGS with Ion AmpliSeq HD laboratory-developed panel (Thermo Fisher Scientific) was used. Analytical validity was ensured by setting the VAF threshold which most accurately identifies primary tumor mutations in plasma cfDNA, whereas clinical validity was evaluated by analyzing HGSOC and low grade serous ovarian cancer (LGSOC) patients since the latter lack *TP53* mutations. To demonstrate that including CTC analysis provides more informative liquid biopsy, compared to ctDNA approach alone, two separate HGSOC cohorts (n=15 each) were enrolled in which for each patient both CTC enriched fraction and plasma cfDNA were analyzed (Figure 1B-D). In cohort 1, CTC enriched fraction was analyzed by gene expression assay, whereas in cohort 2, CTCs were evaluated via *TP53* NGS. The latter was also applied for cfDNA analysis in all 30 patients. Finally, the disease detection rates obtained by CTC and ctDNA assays were compared within each cohort.

**Figure 1.**
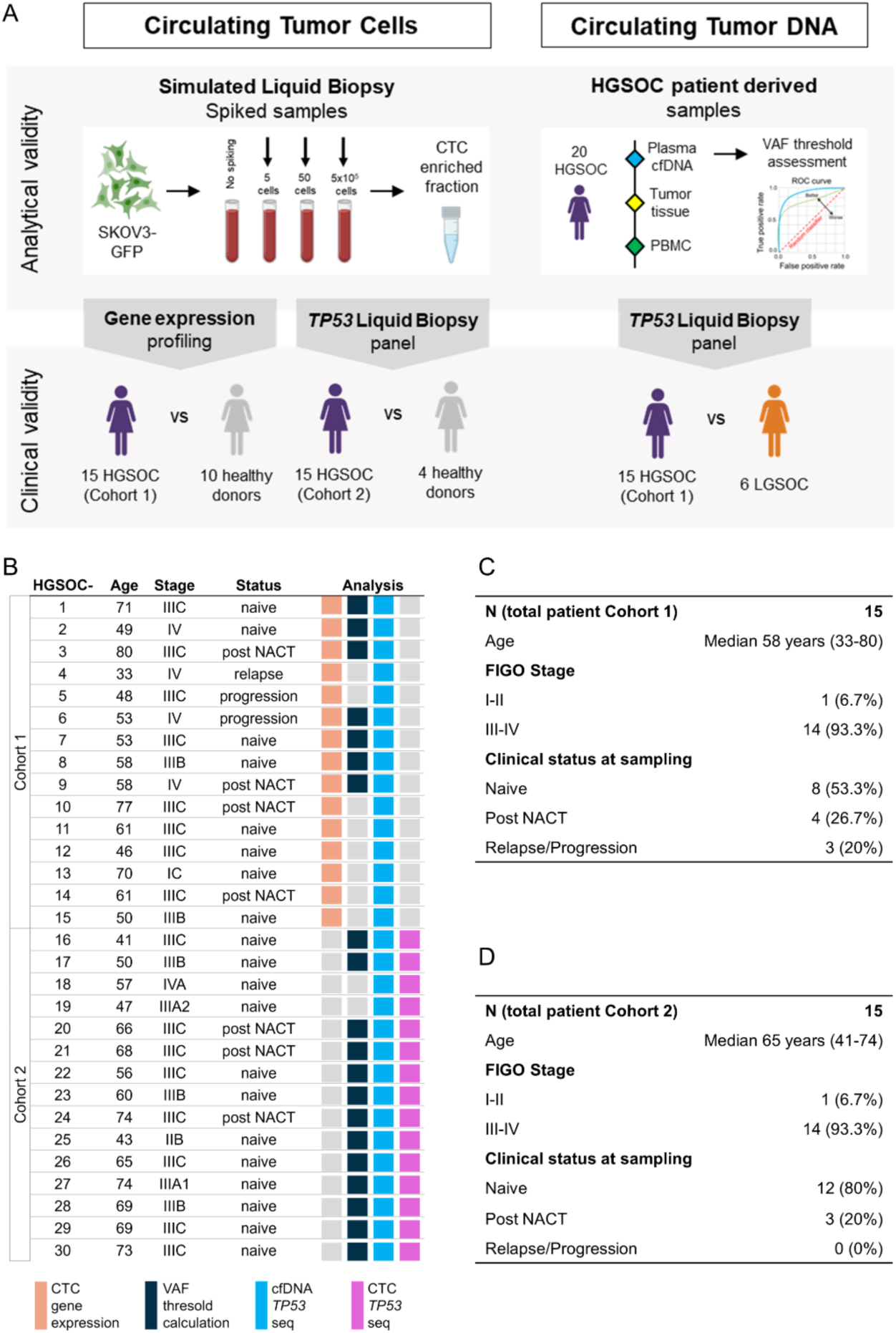
Study design and overview of HGSOC patient characteristics. (FIGO) **(A)** Flowchart reporting the analytical and clinical validity evaluation implemented for CTC and ctDNA detection assays on liquid biopsy samples. **(B)** Demographic and clinical data of HGSOC patients enrolled for the study, with colored squares describing the analyses for which the patient material was used. **(C-D)** Summaries of demographic and clinical data of cohort 1 **(C)** and cohort 2 **(D)** enrolled for the study. International federation of gynecology and obstetrics staging and clinical status are indicated.

### Patients and healthy controls enrolment

(III, IV) For validation of OC CTC gene expression markers, primary tumor tissues of four advanced stage and four early stage (I, II) HGSOC patients who underwent debulking surgery in period between May 2016 and October 2019 were analyzed, together with three PB mononuclear cell (PBMC) samples derived from healthy controls (Supplementary Table 1). Liquid biopsy samples were collected between September 2020 and March 2025, from HGSOC patients who underwent surgery, either primary or interval debulking, at diagnosis or at the disease relapse (n=30, Figure 1B). Inclusion criteria comprised histologic diagnosis of HGSOC, presence of macroscopic disease at surgery in newly diagnosed and relapsed patients, as well as in subjects post neoadjuvant chemotherapy (NACT). Peripheral blood samples used as controls for determining the specificity of CTC analysis via gene expression (n=13), and for analytical validity assessment of CTC detection via *TP53* sequencing (n=4), derived from healthy female donors (inclusion criteria: women aged ≥ 25, with no history of oncologic disease; exclusion criteria: pregnancy, history of oncologic disease) (Supplementary Table 1). Controls for clinical validity assessment of ctDNA *TP53* deep sequencing were patients newly diagnosed with advanced stage LGSOC, whose liquid biopsy was collected between September 2019 and April 2023 (n=6, Supplementary Table 1). *Post-hoc* power calculations were performed using Clincalc [20]. Regarding CTC gene profiling, 88.4% *post hoc* power was calculated to estimate accuracy with alpha of 0.05, considering case *versus* control incidence of 52.5% and 0%, as previously published when using the same method [9]. Since *TP53* sequencing was never used for CTC detection of HGSOC up to date, we considered 66.7% case incidence reported in a breast cancer study which used a mutation-based liquid biopsy assay [21], and 83% *post hoc* power to estimate accuracy with alpha of 0.05 was calculated. Finally, for ctDNA detection, the subjects enrolled for clinical validation allowed 98.8% *post-hoc* statistical power to estimate efficiency, considering 0.05 alpha and case *versus* control incidence of 73% and 0%, respectively [22]. The study was conducted in accordance with the Declaration of Helsinki and the protocol was approved by the Independent Ethics Committee "Comitato Etico di Area Vasta Emilia Centro” (Protocol EM363-2024_107/2011/U/Tess/AOUBo). All subjects enrolled in the study provided written informed consent for their participation.

### Cell lines

OV90 (#ATCC-CRL-3585) and SKOV3 (#ATCC-HTC-77) cells were purchased from ATCC, whereas OC314 (RRID: CVCL_1616) cell line was kindly gifted by Prof. Ada Funaro (University of Turin). Cells were authenticated using AMPFISTRIdentifiler kit (Applied Biosystems #4322288), confirming their putative STR profile. Cells were regularly screened for mycoplasma and were cultured in RPMI 1640 (Euroclone #ECB9006L) supplemented with 10% FBS South America origin EU Approved (Euroclone #ECS5000L), 2 mM L-glutamine (Euroclone #ECB3000D), and 1% penicillin/streptomycin (Euroclone #ECB3001D) in an incubator at 37°C with a humidified atmosphere at 5% CO2. EVOS M5000 Imaging System (ThermoFisher Scientific #AMF5000) was used for cell line monitoring.

### Lentiviral transduction of green fluorescent protein (GFP) in SKOV3 cells

Lentiviral particles expressing GFP were produced using HEK293T cells, seeded in 10 cm tissue culture dishes at 40-50% confluency and incubated overnight. At 80% confluence, cells were transfected using Xtreme Gene Roche Transfection reagent (Cat. No. 6366236001, Roche) according to manufacturer’s instructions. Specifically, the transfection mixture consisted of 7.5 μg psPAX2 packaging plasmid and 2.5 μg pMD2-VSVG envelope plasmid, 10 μg GFP-expressing lentiviral plasmid, (total DNA: 20 μg), diluted in 2 mL Opti-MEM medium. Packaging and envelope plasmids were a kind gift from Prof. Marco Montagner (University of Padova), whereas the GFP-expressing plasmid was a kind gift from Dr. Ilaria Malanchi [23]. Xtreme Gene transfection reagent was added to the DNA with a ratio of 1:3, incubated for 15 min at room temperature, and then added dropwise to the HEK293T cells. Concomitantly, SKOV3 recipient cells were seeded in a T25 flask at 20-30% confluency to achieve optimal cell density for transduction the following day. Viral supernatant was harvested 48 hours post HEK293T transfection and filtered through 0.45 μm syringe filters to remove cellular debris. For SKOV3 transduction, filtered viral particles were mixed 1:1 with fresh culture medium supplemented with polybrene (final concentration: 8 μg/mL) to enhance viral uptake efficiency. SKOV3 cells were incubated with the virus-containing medium for 48 hours. Clonal selection was performed to identify the cells with the highest GFP expression, which was checked with EVOS M5000 Imaging System (Thermo Fisher Scientific #AMF5000), and a pool of 30 clones was made for use in the subsequent experiments.

### Spiked samples preparation

Healthy donor PB (7.5 mL) was collected in 10 mL EDTA tube and manually spiked within 24 hours with known numbers of SKOV3^GFP^ cells: 0, 5, 50, 500, 5’000, or 500’000. In detail, SKOV3 were trypsinized (Sigma #T4049) and counted using Neubauer chamber with Trypan Blue (Sigma #T8154) staining to exclude dead cells. A PBS suspension (200 μL) of 500, 5’000 or 500’000 live cells was then added to PB. To prepare PB samples carrying five, and 50 SKOV3^GFP^ ^s^ingle-cell picking was performed: trypsinized cells were diluted in PBS solution, visualized with phase-contrast microscopy under the sterilized hood, individually picked using 10 μL pipette tip and collected in 1.5 mL Eppendorf tube. PBS was eventually added to final volume of 200 μL and spiked in PB. The same amount of PBS alone was used as a negative control (0 SKOV3^GFP^). Spiked samples were then enriched using Parsortix or CD45 negative sorting for downstream analysis. EVOS M5000 Imaging System (Thermo Fisher Scientific #AMF5000) was used to confirm the presence of SKOV3^GFP^ prior to elution.

### CTC enrichment

At least 7.5 mL of PB was processed for microfluidic-based CTC enrichment by using Parsortix® Cell Separation System (ANGLE) with GEN3D6.5 Cell Separation cassettes according to the manufacturer’s instructions. Default pressure protocol was used, and the captured cells were eluted in 1.2 mL of PBS. For negative affinity-based CTC enrichment, PB was processed using automatic magnetic-activated cell sorting (MACS) for CD45 negative selection. Red Blood Cell Lysis Solution (Miltenyi Bioitec #130-094-183) was used to lyse erythrocytes, and human CD45 MicroBeads (Miltenyi Bioitec #130-045-801) were employed to deplete CD45+ leukocytes, following the manufacturer’s protocol. The stained cell suspension was sorted using AutoMACS® (Miltenyi Biotec) with protocol Depl05 [24]. CTC-enriched solutions obtained from both Parsortix and AutoMACS were centrifuged at 1’200 rpm for 5 min and the pellets were resuspended in 350 μL of RLT buffer for subsequent RNA extraction. For DNA extraction, Parsortix CTC enriched fraction was centrifuged at 1’200 rpm for 5 min and the pellet was stored at -80°C for the subsequent analysis.

### Nucleic acid extraction

RNA extraction from tumor biopsies, cell lines, and CTC-enriched fractions was performed using RNeasy Mini Kit (QIAGEN #74106), while for DNA isolation from formalin fixed paraffin embedded (FFPE) samples Maxwell® CSC DNA FFPE Kit (Promega, #AS1350) was used. Nucleic acid concentrations were assessed with NanoDrop (Thermo Fisher) and stored at -20°C. Peripheral blood for cfDNA analysis was processed within 2 hours from withdrawal and we performed a two-step plasma preparation protocol. In detail, the 10 mL EDTA tube was centrifuged at 3’500 g for 15 min at 4°C, and then plasma was further centrifuged for 10 min at maximum speed. Plasma was stored at -80°C until cfDNA extraction, as we previously confirmed that our preanalytical handling does not affect the downstream analysis. *Quick-*cfDNA Serum&Plasma Kit (Zymo research #D4076) was used for plasma cfDNA extraction, eluted in a final volume of 30 μL. DNA was isolated from CTC enriched fractions by using QIAamp DNA Blood mini kit (QIAGEN #51106) and eluted in a final volume of 30 μL. The same kit was used also for DNA extraction from patients derived PBMC. cfDNA and CTC derived DNA were quantified using Qubit™ (Invitrogen) with High Sensitivity dsDNA assay kit (Invitrogen #Q32851) and stored at -20°C. Liquid biopsy DNA concentration was calculated based on ng/μL eluted after extraction from plasma or CTC-fraction, quantified fluorometrically by using Qubit HS assay, multiplied by μL used and finally divided by either the total mL of plasma analyzed or the total mL of enriched PB.

### cDNA synthesis and preamplification

High-Capacity cDNA Reverse Transcription Kit (Applied Biosystem #4368814) was used for cDNA preparation with random hexamers starting from different RNA concentrations based on samples. When available, 100 ng of RNA was used for cDNA synthesis and we used the maximum possible RNA quantity (14 μL) for CTC-enriched samples, independently from their starting concentration. Reaction was performed using T100 Thermal Cycler (Bio-Rad) following the protocols: 25°C 10 min, 37°C 120 min, 85°C 5 min. A total of 20 μL concentrated cDNA was obtained. For the following preamplification reaction cDNA diluted 1:2 was used as starting material, added with TaqMan® PreAmp Master Mix (2X) (Thermo Fisher Scientific #4384267) and TaqMan gene expression assay of interest pooled at a final concentration of 0.2X, according to the manufacturer’s instructions. Reaction was performed using T100 Thermal Cycler (Bio-Rad) following the protocols: 95°C 10 min; 95°C 15 sec and 60°C 4 min, for 14 cycles, 99°C 10 min. For the downstream gene expression analysis, native cDNA was diluted 1:5, while PA cDNA was diluted 1:2.

### Gene expression quantitative real-time PCR (SybrGreen and TaqMan)

Quantitative RT–PCR was performed using either the intercalating dye SYBR Green dye (Promega) or 5’ nuclease probes PrimeTime™ qPCR Probes (TaqMan assay). For the SYBR Green assay, the primer sequences were designed using Primer3 software [25], the presence of 3’ intra/inter primer homology was excluded using the IDT OligoAnalyzer tool [26] and the availability of the target sequence was estimated by predicting cDNA secondary structure by the Mfold web server [27]. Forward and reverse primers sequences for each target are listed in Supplementary Table 2. For TaqMan assays, the qPCR Probes assay for each gene was selected on the Thermo Fisher website [28], and the predesigned qPCR assays recommended by the manufacturer were used. qRT–PCR with SYBR Green assay was performed with GoTaq qPCR Master Mix (Promega #A6002) and run in 7500 Fast Real-Time PCR System (Applied Biosystem), using the following conditions: 95°C 5 min; 45 cycles of 95°C 15 sec and 63°C 45 sec. The qRT–PCR with TaqMan assay was performed with GoTaq R Probe qPCR Master Mix (Promega #A6101 and #A6102) and run in the abovementioned system, using the following conditions: 95°C 2 min; 40 cycles of 95°C 15 sec and 60°C 1 min.

### Digital PCR

Digital PCR assay was performed using Absolute Q™ DNA Digital PCR Master Mix (5X) (Thermo Fisher Scientific #A52490), TaqMan assay selected for gene expression analysis, and 10 ng of cDNA according to manufacturer’s instructions. The MAP16 plate kit (Thermo Fisher Scientific #A52865) was loaded and run on Absolute Q 22120661 instrument, using the following conditions: 50°C 2 min, 96°C 10 min; 40 cycles of 96°C 5 sec and 60°C 15 sec. The downstream analysis was performed using QuantStudio Absolute Q Software Version 6.3.

### Sequencing of *TP53*

To comprehensively characterize *TP53* genotypes across different sample types, we employed distinct sequencing approaches as follows. Primary tumor tissue was macro dissected from FFPE blocks, DNA was extracted and sequenced using the previously validated NGS laboratory-developed multi-gene panel [29], covering 21.77 kb with 330 amplicons, including *TP53* CDS, as already described in [30]. Mutations were considered valid if present in at least 5% of the total number of analyzed reads and observed in both strands, as *per* previously established validation criteria [31]. The somatic nature of the identified primary tumor mutations was confirmed by Sanger sequencing of matched PBMC following the standard protocol [30]. Liquid biopsy-derived samples (i.e., cfDNA and CTC enriched fraction-derived DNA) were sequenced using the Ion AmpliSeq HD laboratory-developed panel (Thermo Fisher Scientific) targeting the entire coding sequence (CDS) of *TP53* (NM_000546.6, human reference genome hg19/GRCh37), overall covering 1.71 Kb with 39 amplicons. Single-nucleotide variants/Indels present in at least 0.2% of the total number of analyzed reads were considered for variant calling [32–34]. The latter liquid biopsy panel with 0.2% VAF cut-off was also used for sequencing healthy donor control-derived PBMC during analytical validity evaluation for CTC detection, as well as for HGSOC patient-derived PBMC samples sequenced to exclude clonal hematopoiesis as the source of variants found in liquid biopsies. For NGS library preparation, approximately 10 ng of liquid biopsy-derived DNA or 30 ng for FFPE-derived DNA were processed. Sequencing was carried out on an Ion 530 chip. The data were analyzed with the Ion Reporter tool (version 5.20, Thermo Fisher Scientific) and IGV software [35,36] (Integrative Genome Viewer version 2.12.2). Variant classification was performed using American College of Medical Genetics and genomics (ACMG) guidelines with the Varsome database [37,38].

### Statistical analyses

The statistical analyses were performed using R version 4.3.1 with the *pROC*, *dplyr*, *ggplot2*, *binom*, and *presize* packages. Clinical validity was assessed using standard 2x2 contingency tables, which allow calculation of specificity, sensitivity, accuracy, positive predictive value, and negative predictive value. Accuracy estimates of clinical sensitivity and specificity were determined with exact 95% confidence intervals (CI) using the Wilson method. Fisher’s exact test assessed associations between patient group and test results. The Mann-Whitney U test was applied to compare cfDNA VAF between tumor biopsy-specific vs liquid biopsy only variants. For VAF threshold optimization, Receiver Operating Characteristic (ROC) curve analysis was performed to identify the best ctDNA VAF threshold via Youden’s J statistic. The same data set was used for running Leave-One-Out cross-validation (LOOCV) to account for model generalizability, which provided the threshold and relative Area Under the Curve (AUC) for each iteration. ROC statistical significance was evaluated using permutation testing and the Hanley-McNeil Z-test against the null hypothesis of AUC=0.5. Age and cfDNA concentration differences between HGSOC and LGSOC patients were compared using Student’s t-test (GraphPad Prism 8). Statistical significance was defined as p <0.05. Data visualization was performed using GraphPad Prism 8, R version 4.3.1, and BioRender.com.

## Results

### The optimized three-gene expression assay displays high analytical specificity and sensitivity in detecting low cancer cell numbers in simulated liquid biopsies

High inter-patient heterogeneity and ambiguous EpCAM expression in CTCs and PB cells have thus far hampered identification of specific molecular targets for OC CTC detection [39–41]. Previous studies have tried to overcome the issue by relying on techniques which allow multiple marker analysis [6,42,43], the most common strategy being gene expression detection via multiplex qRT-PCR [4,9,10,43,44]. Since there is no consensus on which combination of genes provides the best specificity and sensitivity of such an approach, we first focused on the design and evaluation of an HGSOC gene panel to specifically recognize CTC. The literature search for studies reporting OC CTC gene expression generated a list of 21 targets (Supplementary Table 3), which we validated in a subset of primary tumor tissues deriving from high and low stage HGSOC patients, as well as in PB of three healthy controls (Supplementary Table 1) to account for panel specificity. Most genes (76.2%) were expressed in all tumors, and only *AGR2* resulted to be a less sensitive OC marker, with its expression being observed specifically in high stage HGSOC (Figure 2A). However, nine out of 21 tested genes were also expressed in control samples. This prompted us to test the specificity of the remaining 12 genes in an additional cohort of healthy donor controls (n=10, Supplementary Table 1), by applying TaqMan probe chemistry and cDNA preamplification to increase sensitivity. The results revealed low specificity of *EPCAM*, *ERBB2*, *ERBB3*, *MAL2* and *MUC1*, as these genes were found expressed in 50-80% native cDNA and in 80% of pre-amplified healthy controls (Figure 2B). The analysis of pre-amplified cDNA revealed low specificity also for *PGR, LAMB1, PRAME* and *CDH3*, as they were detected in 30-50% of control samples. On the other hand, *AGR2*, *GPX8* and *WT1* were not expressed in the healthy control cohort and were thus defined as specific HGSOC CTC markers in the context of liquid biopsy (Figure 2B-C).

**Figure 2.**
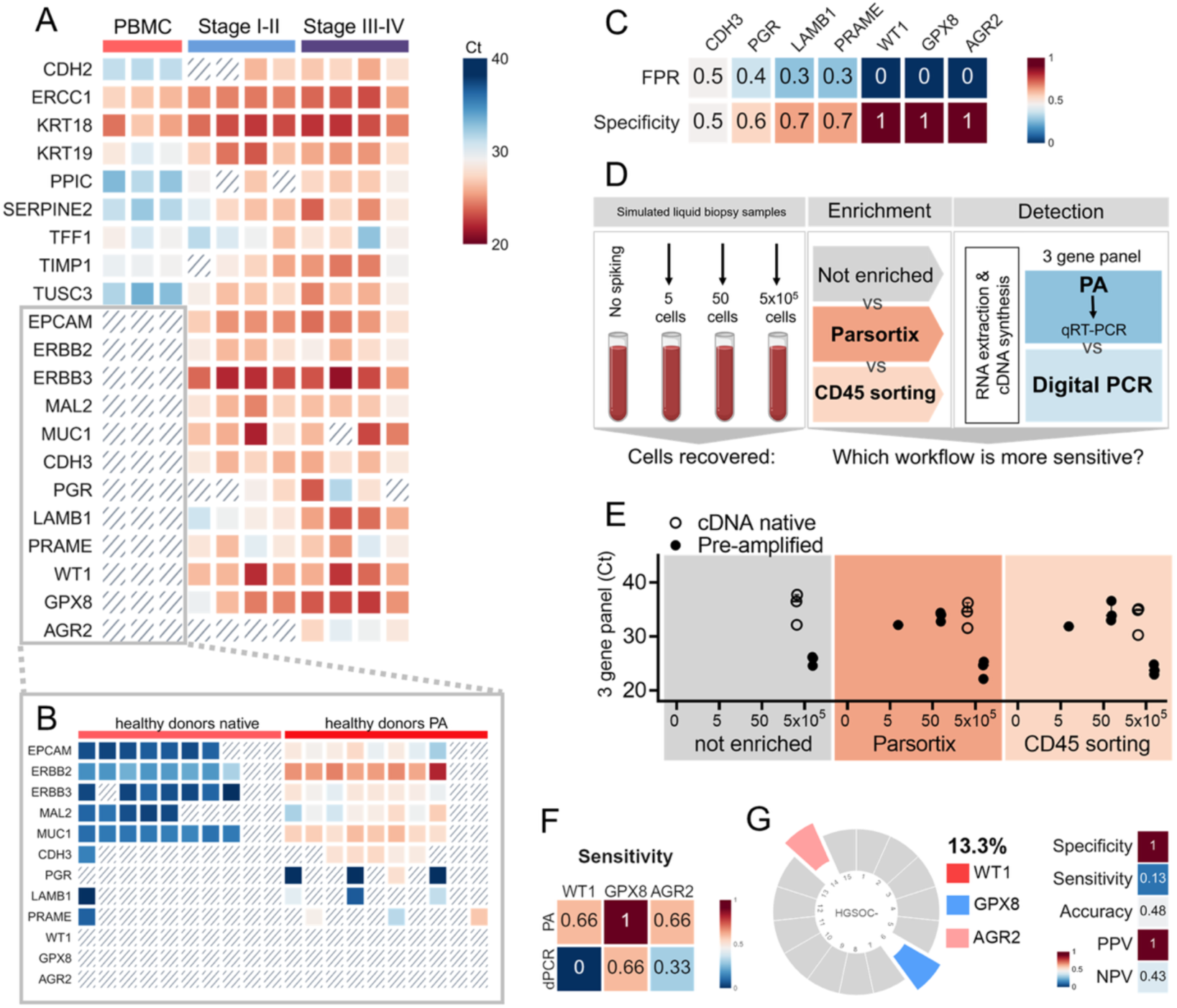
Analytical and clinical validity assessment of the gene expression-based CTC detection in HGSOC patients. **(A)** Cycle-threshold (Ct) heatmaps obtained via qRT-PCR of RNA extracted from three healthy donor PBMCs, eight primary tumors derived from HGSOC patients. Slashed boxes indicate undetected transcripts. **(B)** Cycle-threshold (Ct) heatmaps obtained via qRT-PCR of RNA extracted from PBMCs of ten healthy donors with native or preamplifed (PA) cDNA. Slashed boxes indicate undetected transcripts. **(C)** Heatmap reporting false positive rate (FPR) and specificity values calculated for each gene. **(D)** Graphical sketch of the experimental workflow designed to compare enrichment and detection methods by using simulated liquid biopsy samples. **(E)** Dot plot reporting Cycle-thresholds (Ct) obtained via qRT-PCR of RNA extracted from healthy donor PB spiked with 0, 5, 50, or 500’000 SKOV3 cells. Gray-graph shows not enriched samples. Dark and light orange-graphs show samples enriched with Parsortix and CD45 negative selection, respectively. Native and PA cDNA transcripts are indicated with empty and full circles, respectively. **(F)** Heatmap reporting sensitivity values calculated for each gene by PA cDNA qRT-PCR (PA) and dPCR detection methods. **(G)** Pie chart indicating the disease detection rate of liquid biopsy analysis via Parsortix enrichment coupled with WT1/GPX8/AGR2 CTC detection, and the heatmap reporting the corresponding performance metrics. PPV-positive predictive value; NPV-negative predictive value

Next, we sought to determine which workflow ensures analytical validity for CTC detection via three-gene assay. We used simulated liquid biopsy samples in which known numbers of cancer cells were spiked into healthy donor PB. To choose the optimal model for simulated liquid biopsy preparation, *AGR2*, *GPX8* and *WT1* expression was evaluated in a set of HGSOC cell lines. The highest expression was found in SKOV3 cells (Supplementary Figure 1A), which were then labelled with Green Fluorescent Protein (GFP) to track them during isolation protocols (Supplementary Figure 1B). To mimic the rare cancer cells found in patient liquid biopsies, PB was spiked with five or 50 SKOV3 cells, whereas unspiked samples and spiking 500,000 cells represented negative and positive controls, respectively (Figure 2D). With the secondary aim of comparing CTC enrichment methods, we processed the samples with the two most widely used cancer marker agnostic enrichment approaches, namely Parsortix and CD45 negative magnet assisted cell sorting. The results showed that if samples carrying low cell numbers were not enriched before gene expression analysis, no amplification signals were detected, regardless of the preamplification step (Figure 2E, grey graph). On the other hand, both enrichment protocols allowed detection of even five cancer cells if the cDNA was pre-amplified (Figure 2E, orange graphs), revealing comparable cancer cell recovery efficiency between Parsortix and CD45-negative sorting. Of note, all three genes were detected in samples spiked with 50 cancer cells, whereas only *GPX8* was found in samples carrying five SKOV3 cells (Supplementary Figure 1C). We furthermore investigated whether gene expression analysis via dPCR could provide similar or better sensitivity while avoiding the preamplification step, however such an approach was unable to identify samples spiked with five cells and generally showed lower sensitivity for each of the three genes analyzed (Figure 2F, Supplementary Figure 1D-E).

Taken together, it may be concluded that the three-gene panel analysis, coupled with agnostic enrichment and cDNA preamplification step, is sufficiently sensitive for detection of very low cancer cell numbers, hence tackling the main challenge of CTC-based liquid biopsies.

### Gene expression profiling identifies CTC specific transcripts in a subset of HGSOC liquid biopsies

Once analytical validity was ascertained, we sought to assess the clinical validity of the three-gene panel analysis to detect HGSOC in the CTC-enriched fraction. Regarding the enrichment method, Parsortix was selected as a more suitable procedure due to its marker agnostic property, lower operator hands-on time, and because it potentially captures CTC clusters, otherwise discarded with CD45 negative sorting [11,45,46]. A cohort of 15 HGSOC cases (cohort 1) was enrolled in the study (Figure 1) and liquid biopsies were processed with Parsortix enrichment coupled with the *AGR2*-*GPX8*-*WT1* detection. Only two positive cases were identified, resulting in 0.48 accuracy, 13.3% sensitivity (95% CI: 3.7-37.9 %) and 100% specificity (95% CI: 72.2-100 %) in detecting HGSOC via CTC gene profiling (Fisher’s exact test p=0.5) (Figure 2G). Concordantly with the high heterogeneity of HGSOC, different genes of the panel (*AGR2* and *GPX8*) were found expressed in the two positive patients, HGSOC-6 and HGSOC-14, respectively, suggesting the necessity of a multi-marker approach in increasing performance. Considering the cohort was composed of advanced disease cases, with high macroscopic burden at surgery and high probability of hematogenous spread, it may be concluded that here applied three-gene expression panel is able to detect HGSOC in a low number of patients.

### Optimization of mutation calling ensures analytical validity of *TP53* next-generation sequencing for ctDNA detection in HGSOC liquid biopsy

The clinical value of ctDNA detection in liquid biopsy is being extensively investigated in various cancers, including HGSOC where it is most often performed via sequencing of *TP53* tumor suppressor, which is mutated in virtually all cases [16]. Most often, these analyses have been performed via tumor-informed assays [12–14]. Rare studies dealing with tumor-agnostic cfDNA *TP53* genotyping show promise [15] but call for caution regarding false positive results [47], not infrequent when dealing with low VAF *TP53* variants in plasma cfDNA due to clonal hematopoiesis [48]. Thus, we implemented high-depth *TP53* coding region sequencing of cfDNA via Ion AmpliSeq™ HD high-sensitivity approach, to develop a tumor agnostic assay in which PB ctDNA detection does not rely on previous knowledge about the primary tumor biopsy.

To avoid false positive calls, the first step in the analytical validity assessment was setting the VAF threshold for the mutational call, which guarantees reliable ctDNA detection in liquid biopsy (Figure 3A). Plasma, primary tumor tissue, and PBMC of 20 HGSOC patients were collected and sequenced (Figure 1). We identified *TP53* variants in plasma cfDNA of 17 patients. In total, 31 variants were found, 15 of which (48.39%) matched the primary tumor (Supplementary Figure 2A), presenting median VAF of 1.38% (ranging from 0.2 to 26), and were confirmed to be somatic events upon sequencing corresponding PBMC (Supplementary Figure 2B). Interestingly, the median VAF of the remaining 16 variants was significantly lower (0.27%, ranging from 0.2 to 15.98), suggesting that mutations detected at higher frequencies are more likely to be retrieved in the primary tumor biopsy (Wilcoxon-Mann-Whitney U Test p value=0.006, Figure 3B). To define the most accurate VAF threshold that may discriminate primary tumor driving events from rumor, we performed ROC analysis where solid biopsy confirmed mutations were regarded as true positives and variants absent from matched primary tumor as true negatives. The calculation of Youden’s J factor across different VAFs identified 0.31% as the optimal cut-off, with AUC of 0.77 (SE±0.02, 95% CI 0.58-0.95, Permutation Test p=0.01, Hanley-McNeil Z-test p=0.002) (Figure 3C). Given the limited sample size, we also implemented LOOCV analysis to prevent overfitting. The test confirmed 0.31% as optimal cut-off in 14 out of 17 iterations (82.35%, mean VAF 0.33%±0.05 SD) and revealed AUC values ranging from 0.75 to 0.83 (mean training AUC 0.77; Figure 3D), demonstrating consistent discriminative ability of the model.

**Figure 3.**
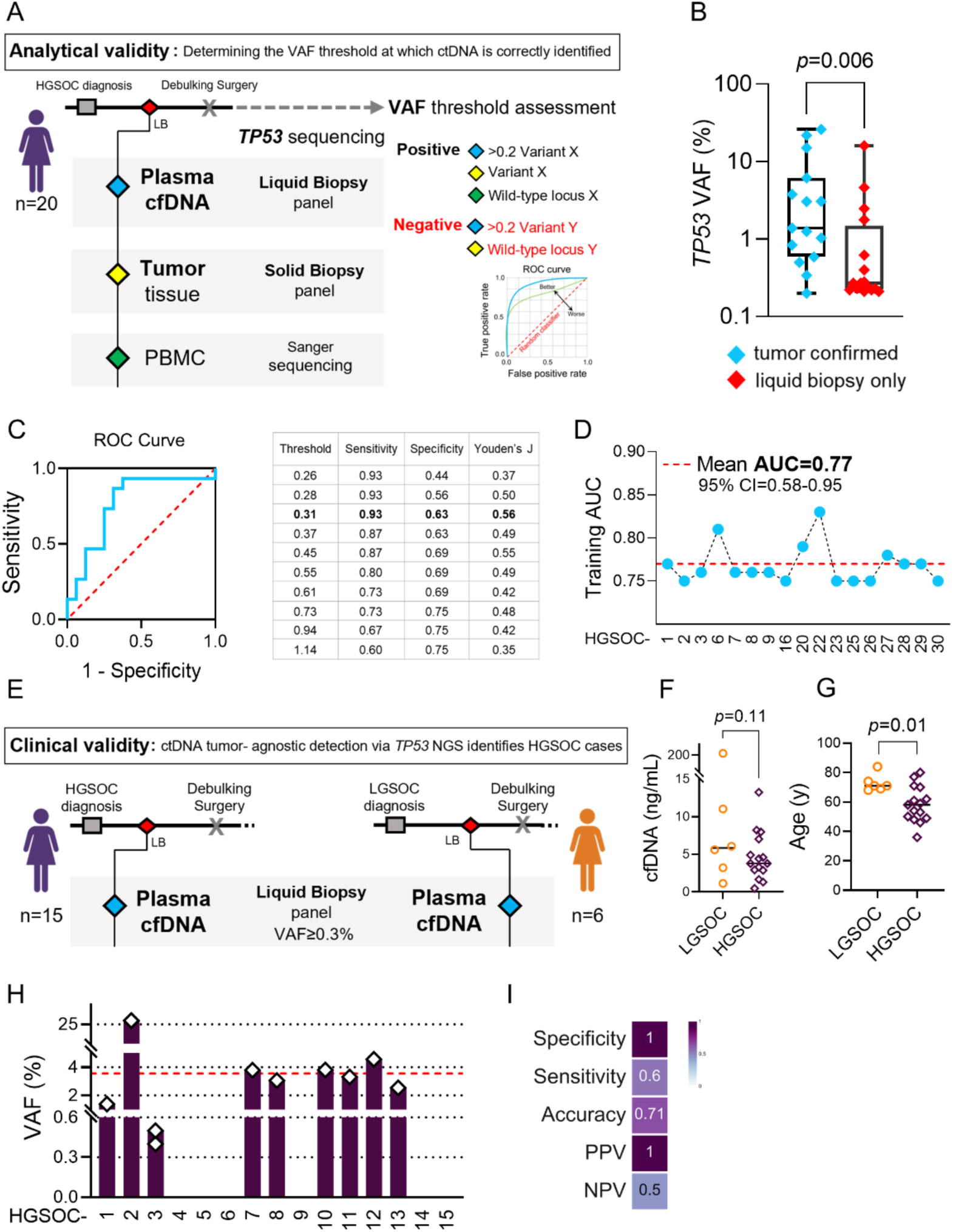
Analytical and clinical validity assessment of tumor-agnostic TP53 NGS-based ctDNA detection in HGSOC. **(A)** Graphical sketch of the workflow for definition of ctDNA detection VAF cut-off, performed by sequencing cfDNA variants in matched primary tumor tissue and PBMC. **(B)** Primary tumor confirmed (blue) and cfDNA private (red) TP53 VAFs found in HGSOC patients’ plasma. P value indicates Mann-Whitney U test result. **(C)** ROC curve (blue) computed by considering tumor confirmed and cfDNA private TP53 VAFs as positive and negative category, respectively. Dashed red line indicates the random classifier. Adjacent table lists the sensitivity, specificity, and Youden’s J calculated for each VAF threshold. The optimal cut-off value is indicated in bold. **(D)** Dot-plot reporting the LOOCV analysis training AUC results. The X axis reports HGSOC patient used to test the VAF threshold obtained from each iteration. Red dashed line indicates the mean AUC. **(E**) Graphical sketch of the workflow for clinical validity assessment. HGSOC and LGSOC liquid biopsies (LB) were used as positive and negative controls, respectively. **(F)** Median plasma cfDNA concentration in LGSOC (orange) and HGSOC (purple) patients. Single dots represent patients, black bars the computed median values, and p value indicates the t-test result. **(G)** Age in years (y) at liquid biopsy sampling of LGSOC (orange) and HGSOC (purple) patients. Single dots represent patients, black bars the computed median values, and p value indicates the t-test result. **(H)** TP53 VAFs found in plasma cfDNA of HGSOC cohort 1. Dashed red line indicates the median value. **(I)** Performance metrics heatmap of the tumor-agnostic TP53 ctDNA detection assay. PPV-positive predictive value; NPV-negative predictive value.

Taken together, our optimization of the mutation calling provides evidence for analytical validity of high-depth *TP53* sequencing for ctDNA detection in HGSOC liquid biopsies.

### Customized *TP53* deep sequencing is a reliable approach for liquid biopsy-based detection of HGSOC

With the aim of measuring the performance of the test to reliably and reproducibly detect HGSOC, the clinical validity was next assessed for the *TP53* NGS-based ctDNA assay (Figure 3E). Fifteen HGSOC patients from cohort 1 were considered as positive cases. As negative controls, liquid biopsies derived from six LGSOC patients were collected (Supplementary Table 1), since they mimic tumor burden and corresponding high cfDNA concentration but lack *TP53* mutations [49]. The LGSOC cohort showed comparable cfDNA concentrations (Figure 3F) but was significantly older than the high-grade counterpart (Figure 3G). Since potential false positives due to clonal hematopoiesis are expected to occur more likely in elderly subjects, the age difference was not expected to bias the clinical validity assessment.

Collectively, considering the defined 0.31% VAF threshold, all LGSOC derived cfDNA were *TP53* wild-type, indicating 100% specificity of the assay. On the other hand, in the HGSOC cohort, 10 *TP53* variants were identified in nine patients, with a median VAF of 3.55%, ranging from 0.4% to 26% (Table 1, Figure 3H), revealing 0.71 accuracy, 60% sensitivity (95% CI: 37.5-80.2%) and 100% specificity (95% CI: 61-100%) in detecting HGSOC via ctDNA approach (Fisher’s exact test p=0.01) (Figure 3I).

**Table 1.** TP53 variants and their allele frequencies found by cfDNA sequencing in liquid biopsies of cohort 1. del=deletion; fs=frameshift.

| HGSOC- | ctDNA VAF (%) | *TP53* variant |
| --- | --- | --- |
| 1 | 1.38 | c.376-2A>G |
| 2 | 26 | p.T253_I254del |
| 3 | 0.5 | p.Q167Hfs*3 |
|  | 0.4 | p.Y205C |
| 4 | 0 | / |
| 5 | 0 | / |
| 6 | 0 | / |
| 7 | 3.78 | p.G245D |
| 8 | 3.05 | p.L201Ffs*8 |
| 9 | 0 | / |
| 10 | 3.8 | p.E287Rfs*58 |
| 11 | 3.31 | p.I195T |
| 12 | 4.56 | p.M246T |
| 13 | 2.53 | p.Y126* |
| 14 | 0 | / |
| 15 | 0 | / |

Taken together, our results establish clinical validity of cfDNA *TP53* sequencing as a sensitive and specific approach for HGSOC detection via liquid biopsy and suggest 0.31% VAF threshold for ctDNA mutation calling when the Ion AmpliSeq™ HD system is used.

### Simultaneous application of CTC and ctDNA identification assays improves HGSOC detection via liquid biopsy

The combined detection of CTC and ctDNA has demonstrated additional clinical value over single-analyte approaches in various cancer types [21,50–52]. However, this dual strategy has not yet been explored in the context of HGSOC, where it may offer substantial benefits to patients if shown to improve the rate of disease detection. Therefore, we compared the data obtained by the two assays in cohort 1 (Figure 4A), in which the presence of CTCs was evaluated by gene expression. Four cases displayed concordant negativity (26.7%), whereas no cases resulted positive for both assays. Interestingly, the two patients positive for CTCs were concomitantly negative for ctDNA (13.3%), and nine patients positive for ctDNA were negative for CTCs (60%), showing that the combinatory approach indeed increased HGSOC detection rate compared to either single analyte assay (Figure 4B). Collectively, these results suggest that including CTC identification in the workflow increases the disease detection rate from 60%, obtained by the cfDNA-based analysis, to 73.3%, showing potential for improving the clinical utility of liquid biopsy for HGSOC.

**Figure 4.**
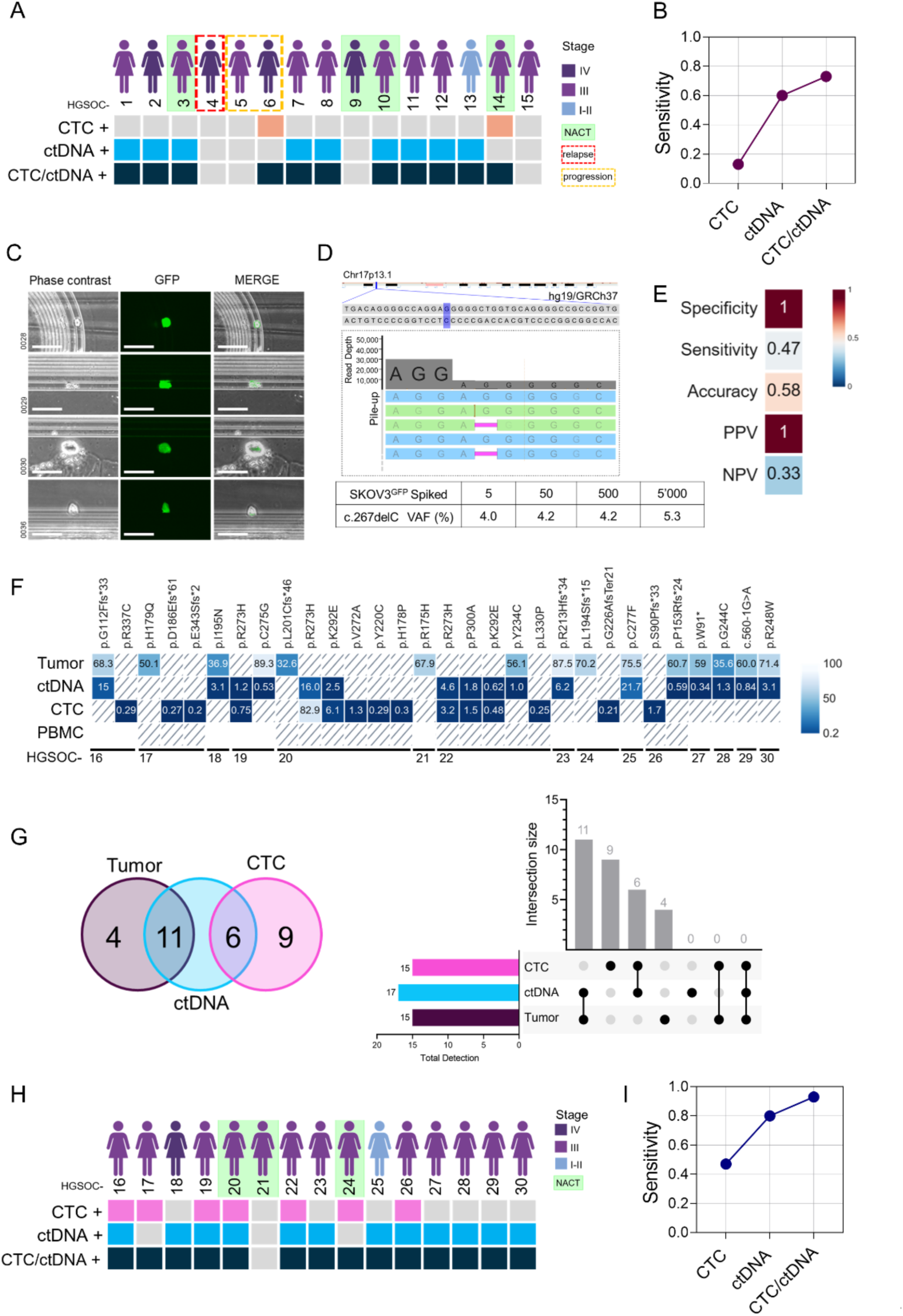
Combining CTC and ctDNA detection in HGSOC liquid biopsies. **(A)** Schematic representation of CTC and ctDNA detection results across cohort 1. Colored and gray squares indicate positive and negative results, respectively. Tumor stage and clinical status are indicated. **(B)** Sensitivity of CTC, ctDNA, and combined detection assays across cohort 1. **(C)** Images of four SKOV3^GFP^ captured by Parsortix cassette while processing healthy donor PB spiked with five cells. Scale bar = 100 μm. **(D)** Representative GenomeBrowser output image of the TP53 NGS of the sample spiked with five SKOV3 cells. The pink bar indicates c.267delC mutation locus, with blue and green bars indicating complementary sequencing reads. Read depth is presented on y-axis. The adjacent table lists TP53 c.267delC VAF found in all spiked samples. **(E)** Heatmap reporting the performance metrics of tumor-agnostic TP53 sequencing for CTC detection. PPV-positive predictive value; NPV-negative predictive value. **(F)** VAF heatmap of TP53 mutations identified in solid and liquid biopsies of cohort 2. Slashed boxes indicate wild-type genotype, whereas white boxes represent samples which were not available for the analysis. **(G)** Venn diagram and UpSet plot showing shared TP53 variants across tumor tissue, ctDNA and CTC samples. The horizontal bar plot indicates the number of variants found in each analyte, whereas the vertical bar plot shows the number of variants shared between various sample types (Intersection size), according to the black dot intersections found below. **(H)** Schematic representation of CTC and ctDNA detection results across cohort 2. Colored and gray squares indicate positive and negative results, respectively. Tumor stage and clinical status are indicated. **(I)** Sensitivity of CTC, ctDNA, and combined detection assays across cohort 2.

Driven by these results, we next sought to corroborate the finding in a second HGSOC cohort and by applying a different CTC detection method. To harmonize the workflow and reduce turnaround times of tandem CTC/ctDNA analysis, *TP53* liquid biopsy panel was chosen to detect both analytes. To determine whether the approach provides high sensitivity and specificity for CTC detection, analytical validity assessment was performed via *TP53* sequencing of the Parsortix-enriched simulated liquid biopsies, prepared by spiking 5, 50, 500 or 5’000 SKOV3^GFP^ in healthy donor PB. Upon Parsortix enrichment, SKOV3^GFP^ were detected in all spiked samples, including the one carrying five cancer cells where four events were observed (Figure 4C). Subsequently, high depth *TP53* sequencing was performed and the SKOV3 specific c.267delC mutation was detected in all spiked samples (Figure 4D), demonstrating high analytical sensitivity of the approach. No other mutations were identified, suggesting also high analytical specificity, which was furthermore confirmed by analysing four age-matched healthy donors’ PBMC (Supplementary Table 1), in which no *TP53* variants were detected (https://www.ncbi.nlm.nih.gov/bioproject/PRJNA1338131). Once analytical validity of deep *TP53* sequencing for both analytes was ensured, plasma cfDNA and Parsortix-enriched corpuscular fraction were sequenced in 15 HGSOC patients (cohort 2, Figure 1B and Figure 1D), to compare CTC and ctDNA-based liquid biopsy performances. In the CTC fraction, 0.58 accuracy, 46.7% sensitivity (95% CI: 24.8-69.9%), and 100% specificity (95% CI: 43.9-100%) were achieved (Fisher’s exact test p=0.24) (Figure 4E). Specifically, 15 variants in seven patients were detected, with 0.48% median VAF, ranging from 0.21 to 82.9 (Figure 4F). On the other hand, cfDNA sequencing, similarly to results obtained in cohort 1, detected 17 variants in 12 patients, with median VAF of 1.77% (ranging from 0.34 to 21.73) (Figure 4F), and 80% disease detection rate, establishing the solid performance of the assay.

To compare genetic heterogeneity between liquid and solid biopsy variants, we retrieved primary tumor TP53 genotyping data which revealed that 35.3% of ctDNA variants were undetected in the solid biopsy (Figure 4F-G). Moreover, variants found in CTC fraction were either private (60%) or shared with ctDNA (40%) (Figure 4F-G). We ruled out the possibility of these being a consequence of clonal haematopoiesis since no TP53 alterations were detected by high-depth sequencing in matching PBMC of patients whose material was available (Figure 4F, https://www.ncbi.nlm.nih.gov/bioproject/PRJNA1338131). The latter result, together with the efficient analytical validity assessment, where no unspecific variants were detected in control samples, allowed to reasonably define ctDNA/CTC private variants as true HGSOC-associated events, probably explained by tumor clonal dynamics [53].

We next compared the disease detection rate obtained by the two assays in cohort 2 (Figure 4H). One case displayed concordant negativity (6.7%), whereas five patients were positive by both assays (33.3%). Finally, two patients positive for CTCs were concomitantly negative for ctDNA (13.3%) and seven patients positive for ctDNA were negative for CTCs (46.7%), again showing that the combinatory approach increases the ability to detect HGSOC. Specifically, including both analyses in the liquid biopsy protocol increased the sensitivity to 93.3% (Figure 4I).

Taken together, ctDNA detection resulted to be more sensitive and more accurate liquid biopsy approach for HGSOC detection compared to CTC-based assays here applied. Provided adequate optimization of CTC identification is achieved, our data imply that tandem CTC/ctDNA analysis may improve clinical utility of liquid biopsy in HGSOC.

## Discussion

Liquid biopsy approaches based on the detection of CTCs or ctDNA have been extensively investigated across a wide range of malignancies for both early cancer detection and prediction of therapeutic response. In certain clinical contexts, these methodologies have already been integrated into routine practice [1], but not in HGSOC, where no screening protocols are currently available, leading most-often to late-stage diagnosis, with consequent high mortality and complicated patient management. Our study represents a step forward in advancing liquid biopsy strategies for HGSOC, demonstrating that while cfDNA analysis outperforms CTC-based assays for disease detection, the combined approach significantly enhances clinical sensitivity compared to either analyte alone. These results highlight the potential translational value of the simultaneous ctDNA and CTCs in HGSOC.

It is important to emphasize that, to establish the proof of principle for increased clinical sensitivity of the tandem CTC/ctDNA analysis, a rigorous study design was implemented, ensuring high analytical specificity and sensitivity of each assay prior to clinical validity assessment. In the context of CTC analysis, the three-gene expression assay and *TP53* deep sequencing, both combined with Parsortix enrichment, demonstrated the capability to detect as few as five cancer cells spiked into PB. This level of sensitivity is comparable to that reported for other available methodologies, such as the immunocapture-based AdnaTest Ovarian Cancer Detect [43] and CellSearch [54]. A key strength of our approach was assessing assay specificity by including healthy donor samples as negative controls, which are often omitted in similar studies [43,54], or yield false positive results when included [44]. For ctDNA detection, we employed a laboratory-developed *TP53* sequencing panel coupled to the Ion AmpliSeq HD system. Analytical validity was ensured by empirically identifying 0.31% as the optimal VAF threshold for accurate mutation detection. Given that the reliable calling of low-frequency ctDNA variants (<0.5% VAF) is widely recognized as a critical challenge in the field [33], our established limit of detection represents a meaningful advancement in minimizing false-negative results. Notably, while some studies exclude variants with high allele frequencies to avoid inclusion of germline mutations [55], our data demonstrated that tumor-specific ctDNA variants were frequently present above >1% threshold, highlighting the limitations of such exclusion criteria.

Regarding clinical validity, the performance comparison between cfDNA and CTC-based assays revealed the former as a more efficient liquid biopsy strategy. While both showed 100% specificity, cfDNA assay resulted in higher clinical sensitivity, which was either similar (cohort 1; 60%) or slightly superior (cohort 2; 80%) to those of previous assays based on *TP53* cfDNA sequencing in HGSOC, reporting clinical sensitivity of 60–73% [22,56,57]. On the other hand, studies evaluating CTCs in OC display a wide range of detection rates, from 15% to 98% [58– 63], reflecting substantial methodological heterogeneity. Our findings align with studies reporting lower sensitivity of CTC-based assays, which may be attributed to the stringent specificity criteria defined in our study design. The latter were deliberately selected based on recent evidence indicating that less rigorous methods for capturing OC CTCs are associated with an increased risk of false-positive results [41]. Indeed, during validation of the three-gene expression assay, we observed unexpectedly high expression levels of two keratin-encoding genes in healthy controls, biomarkers frequently used in oncologic liquid biopsy assays [11]. This observation underlines the need for caution when incorporating such markers in assays for HGSOC CTC detection. Another explanation for low disease detection rate observed with CTC-based assays could be the technological constraints intrinsic in our approach. We have based our choice of genetic CTC detection with the aim to develop an assay with reasonable turnaround times which could easily be implemented in molecular pathology labs. With hindsight we believe CTC enumeration techniques may provide better performance than here applied genetic methods and should, thus, be considered in future dual-analyte studies.

Interestingly, *TP53* sequencing resulted superior compared to the three-gene detection assay for CTC-based HGSOC identification. However, it must be acknowledged that CTC variants were undetected in the matching primary tumor, similarly to more than a third of ctDNA variants found in cohort 2. We excluded this might be due to sequencing errors, since the same high-depth NGS approach revealed no *TP53* variants in healthy donor controls, spiked samples (with exception of the SKOV3 variant) or LGSOC liquid biopsies. Moreover, we excluded the possibility of these being a consequence of clonal hematopoiesis, by sequencing patients’ matching PBMCs, allowing to reasonably consider these liquid biopsy private mutations as HGSOC-associated events, possibly explained by clonal dynamics [53], selective immunoclearance of extravasated CTCs [64] or patient cancerous field effect [65]. Such a conclusion is furthermore supported by the fact that all ctDNA variants undetected in primary tissue (35.3%) were shared with those identified in CTCs. In line with our findings, several studies report liquid-biopsy private variants [50–52,66], and clinically relevant tumor genetic heterogeneity has also been observed in HGSOC cfDNA [67,68]. The peculiarity of our finding is that *TP53* mutations are usually considered clonal HGSOC markers [67]. To the best of our knowledge, only a few recent studies, by applying 20,000-30,000 sequencing depth, comparable to here applied NGS approach, find cfDNA private *TP53* variants for which they also exclude clonal hematopoiesis derivation and hypothesize tumor origin [15,69].

Importantly, the fact that tumor undetected *TP53* variants where often shared between ctDNA and matched CTC fraction, implies that analyzing both analytes together enables mutation cross-validation, which could permit to avoid invasive tumor biopsy in the first-line blood-based HGSOC screening.

Our most relevant finding regards the substantial increase in disease detection rate observed when results of both CTC and ctDNA assays were combined, proving such an approach allows a more efficient liquid biopsy. This is concordant with studies combining CTC and cfDNA analysis in other cancers, which generally agree on the consideration that both analytes broaden the applicability of liquid biopsy in oncologic patients [21,50,66]. Interestingly, Alba-Bernal et al. reveal that, while cfDNA analysis is more efficient in detecting pre-treatment breast tumors, CTCs are found more often than ctDNA in matching post-treatment and post-surgery liquid biopsies, suggesting their particular value when assessing therapy response and minimal residual disease (MDR) [21]. Moreover, similarly to our discovery of private *TP53* variants in CTC fraction, other studies have shown higher genetic heterogeneity of CTCs compared to ctDNA [21], meaning that their investigation could provide complementary data useful for patient management in terms of therapeutic choice. In this context, extending mutation screening to other tumor suppressors and oncogenes associated with HGSOC might discover broader genetic heterogeneity and offer even more informative liquid biopsy.

While demonstrating, for the first time, that tandem CTC/ctDNA analysis improves HGSOC liquid biopsy, a limitation remains, which stands in the relatively small cohort of subjects analysed. This prevented us from drawing conclusions on correlation between liquid biopsy results and clinical parameters, such as progression free/overall survival, CA-125 biomarker values or radiology findings. Indeed, the clinical utility of the approach is still to be evaluated, by including a higher proportion of stage I/II cases, as well as longitudinally collected liquid biopsies during the patient follow-up, to allow evaluation of the tandem analysis efficiency for early detection, therapy response prediction and MRD assessment.

## Conclusion

Overall, our findings advance liquid biopsy strategies in HGSOC by showing that, while cfDNA outperforms CTCs alone, their combination improves the disease detection rate. Whereas ctDNA provides a sensitive snapshot of tumor burden and enables real-time monitoring of disease dynamics, CTCs reflect historic clonal heterogeneity of the malignancy. The complementary nature of these biomarkers suggests that a multi-analytic approach holds translational potential for integrating liquid biopsy into precision oncology programs of HGSOC, to possibly allow earlier diagnosis, relapse detection, better patient stratification, and anticipate treatment resistance before radiologic progression.

## List of abbreviations

(ACMG): American College of Medical Genetics and Genomics
(AUC): Area Under the Curve
(cfDNA): Circulating cell-free DNA
(CTCs): Circulating tumor cells
(ctDNA): Circulating tumor DNA
(CI): Confidence interval
(dPCR): Digital PCR
(EpCAM): Epithelial cell adhesion molecule
(FFPE): Formalin fixed paraffin embedded
(GFP): Green fluorescent protein
(HGSOC): High grade serous tubo-ovarian carcinoma
(LOOCV): Leave-One-Out cross-validation
(LGSOC): Low grade serous ovarian cancer
(MACS): Magnetic-activated cell sorting
(NACT): Neoadjuvant chemotherapy
(NGS): Next-generation sequencingOvarian cancer (OC)
(PB): Peripheral blood
(PBMC): Peripheral blood mononuclear cell
(qRT-PCR): Quantitative Real Time-PCR
(ROC): Receiver Operating Characteristic
(VAF): Variant allele frequencies

## Declarations

### Ethics approval and consent to participate

This study was conducted in accordance with the Declaration of Helsinki and the protocol was approved by the Independent Ethics Committee "Comitato Etico di Area Vasta Emilia Centro” (Protocol EM363-2024_107/2011/U/Tess/AOUBo). All subjects enrolled in the study provided written informed consent for their participation.

### Consent for publication

Not applicable

### Availability of data and materials

All data generated and analysed during this study are included in this published article and its supplementary information file, apart from raw device output data, which are deposited in https://www.ncbi.nlm.nih.gov/bioproject/PRJNA1338131 and https://doi.org/10.6092/unibo/amsacta/8538.

## Authors’ contributions

Conceptualization, B.C., I.K.; methodology, B.C., D.d.B.; acquisition and analysis, B.C., S.Cor., M.D.L., S.L., C.A.C., G. Gi., S. Col., A.D.L., D.d.B., I.K.; data interpretation: B.C., S.d.C., F.M., P.D.I., A.Ma.P., A.My.P., D.d.B., G.Ga., I.K.; writing – original draft, B.C., I.K., writing – review & editing, B.C., C.A.C, A.Ma.P., A.My.P., D.d.B., G.Ga, I.K.; project administration, I.K., funding acquisition and resources, A.Ma.P., G.Ga., I.K. All authors have approved the submitted version of the work and have agreed both to be personally accountable for the author’s own contributions and to ensure that questions related to the accuracy or integrity of any part of the work, even ones in which the author was not personally involved, are appropriately investigated, resolved, and the resolution documented in the literature.

## Acknowledgements

Not applicable

