## Supplementary Material for "Combining circulating tumor cell and circulating cell free DNA analyses enhances liquid biopsy sensitivity in detecting high grade serous tubo-ovarian carcinoma"

Cavina *et al.*

**Table of contents:**

Supplementary Tables 2

Supplementary Figures 5

Supplementary References 7

### Supplementary Tables

***Supplementary Table 1. Demographic and clinical characteristics of participants used for assay validations:*** *HGSOC patients (T1-T8) and healthy donor controls (CNTR1-3) whose tumor tissue and PBMC samples, respectively, were used for target gene validation in Figure 2A; Healthy donor controls (CNTR4-13) whose PBMC samples were used for target gene specificity validation in Figure 2B; Healthy donor controls (CNTR14-16) whose PBMC samples were used for specificity assessment of high-depth TP53 sequencing assay for CTC detection; LGSOC patients (L1-L6) whose plasma cfDNA was used for specificity assessment of high-depth TP53 sequencing assay for ctDNA detection.*

| ID | AGE | FIGO stage | Analysis |
|---|---|---|---|
| T1 | 66 | IV | Gene validation cases and controls (Figure 2A) |
| T2 | 39 | IV | |
| T3 | 69 | IIIC | |
| T4 | 60 | IIIC | |
| T5 | 68 | IC | |
| T6 | 65 | II | |
| T7 | 70 | IC | |
| T8 | 54 | II | |
| CNTR1 | 40 | / | |
| CNTR2 | 44 | / | |
| CNTR3 | 25 | / | |
| CNTR4 | 44 | / | Gene specificity validation controls (Figure 2B) |
| CNTR5 | 40 | / | |
| CNTR6 | 28 | / | |
| CNTR7 | 26 | / | |
| CNTR8 | 25 | / | |
| CNTR9 | 43 | / | |
| CNTR10 | 39 | / | |
| CNTR11 | 25 | / | |
| CNTR12 | 52 | / | |
| CNTR13 | 64 | / | |
| CNTR14 | 74 | / | Controls for *TP53* sequencing assay (CTC detection) |
| CNTR15 | 69 | / | |
| CNTR16 | 73 | / | |
| CNTR17 | 69 | / | |
| L1 | 74 | IIIC | Controls for *TP53* sequencing assay (ctDNA detection) |
| L2 | 84 | IIIC | |
| L3 | 71 | IIIC | |
| L4 | 68 | IIIB | |
| L5 | 69 | IV | |
| L6 | 71 | IIIC | |

***Supplementary Table 2. Target gene primer sequences used for SYBR qRT-PCR analysis.***

| Gene | Forward | Reverse |
|---|---|---|
| *PGR* | ACCTGACACCTCCAGTTCTT | TCCATCCTAGACCAAACACCA |
| *CDH3* | GAAGCGGAAGATCAAGGAGC | GTGGAGCTGGGTGATGTCAT |
| *CDH2* | GGCTTCTGGTGAAATCGCAT | TCCACCTTAAAATCTGCAGGC |
| *TUSC3* | AAGCAGCAACTTCGAAAGGC | GGATAGCCGTGGTACTTGGA |
| *MAL2* | GCCACATCCCTGCATGATTT | CGGTCGCCATCTTCGTAAAG |
| *LAMB1* | GAAGCTTTCGGTGACCTCG | CTGTCAGGATTCAGGGTCTCA |
| *SERPINE2* | CCTCTGCCTGTGATTCCATC | CTTTGTGTTCTCGGGTTGGA |
| *PRAME* | CTGCAGGCTCTCTATGTGGA | GACACTTAGCTGACTGACGC |
| *AGR2* | ATTGGCAGAGCAGTTTGTCC | TCTTCCAGTGATATCGGCTCT |
| *MUC1* | CCAGTCTCCTTTCTTCCTGC | TTCTCTGGGTAGCCGAAGTC |
| *GPX8* | CGGAGTAACTTTCCCCATCTTC | TGGCTTCCAGAACTTCACAAC |
| *EPCAM* | ACTACAAGCTGGCCGTAAAC | CCCCTTCAGGTTTTGCTCTTC |
| *KRT18* | CTTGGAGAAGAAGGGACCCC | GTCATCAGCAGCAAGACGG |
| *KRT19* | GAGCAGGTCCGAGGTTACTG | CTGGGCTTCAATACCGCTG |
| *ERCC1* | CTGGGAAGGACAAAGAGGGG | ATTCGGCGTAGGTCTGAGG |
| *PPIC* | GCAGAATTGTGATTGGCCTC | TGATGACACGATGAAACTTGCT |
| *WT1* | AGAGCGATAACCACACAACG | CATGAAGGGGCGTTTCTCAC |
| *TFF1* | AGACGTGTACAGTGGCCC | GGAGGGACGTCGATGGTATT |
| *ERBB3* | GACAACCTGGCAACCAC | GCTCCCAGAAACTGCAGACT |
| *TIMP1* | TACTTCCACAGGTCCCACAA | ACAGCCAACAGTGTAGGTCT |
| *ERBB2* | ACCTGGAACTCACCTACCTG | TTGTCCTCAAAGAGCTGGGT |

***Supplementary Table 3. NCBI genes selected as putative ovarian cancer (OC) markers from scientific literature (Ref).*** *Na-Not available.*

| Gene ID | Symbol | Full name | Ref. | Number of patients | Positivity rate (%) |
|---|---|---|---|---|---|
| 5241 | *PGR* | progesterone receptor | [1] | 19 | 0 |
| 102886151 | *CDH3* | cadherin 3 | [2] | 200 | 4 |
| 1000 | *CDH2* | cadherin 2 | [1] | 19 | 36.8 |
| 7991 | *TUSC3* | tumor suppressor candidate 3 | [2] | 200 | 3.5 |
| 114569 | *MAL2* | mal, T cell differentiation protein 2 | [1] | 19 | 31.6 |
| 3912 | *LAMB1* | laminin subunit beta 1 | [2] | 200 | 1.5 |
| 5270 | *SERPINE2* | serpin family E member 2 | [1] | 19 | 89.5 |
| 23532 | *PRAME* | PRAME nuclear receptor transcriptional regulator | [1] | 19 | 15.8 |
| 10551 | *AGR2* | anterior gradient 2, protein disulphide isomerase family member | [2] | 200 | 0.5 |
| 4582 | *MUC1* | mucin 1, cell surface associated | [3] | 137 | Na |
| 493869 | *GPX8* | glutathione peroxidase 8 (putative) | [2] | 200 | 5.5 |
| 4072 | *EPCAM* | epithelial cell adhesion molecule | [2] | 200 | 2 |
| 3875 | *KRT18* | keratin 18 | [4] | 40 | Na |
| 3880 | *KRT19* | keratin 19 | [4] | 40 | Na |
| 2067 | *ERCC1* | ERCC excision repair 1, endonuclease non-catalytic subunit | [5] | 143 | 8 |
| 5480 | *PPIC* | peptidylprolyl isomerase C | [2] | 200 | 17 |
| 7490 | *WT1* | WT1 transcription factor | [4] | 40 | Na |
| 7031 | *TFF1* | trefoil factor 1 | [2] | 200 | 0 |
| 2065 | *ERBB3* | erb-b2 receptor tyrosine kinase 3 | [1] | 19 | 26.3 |
| 7076 | *TIMP1* | TIMP metallopeptidase inhibitor 1 | [6] | 38 | Na |
| 2064 | *ERBB2* | erb-b2 receptor tyrosine kinase 2 | [1] | 19 | 84.2 |

### Supplementary Figures

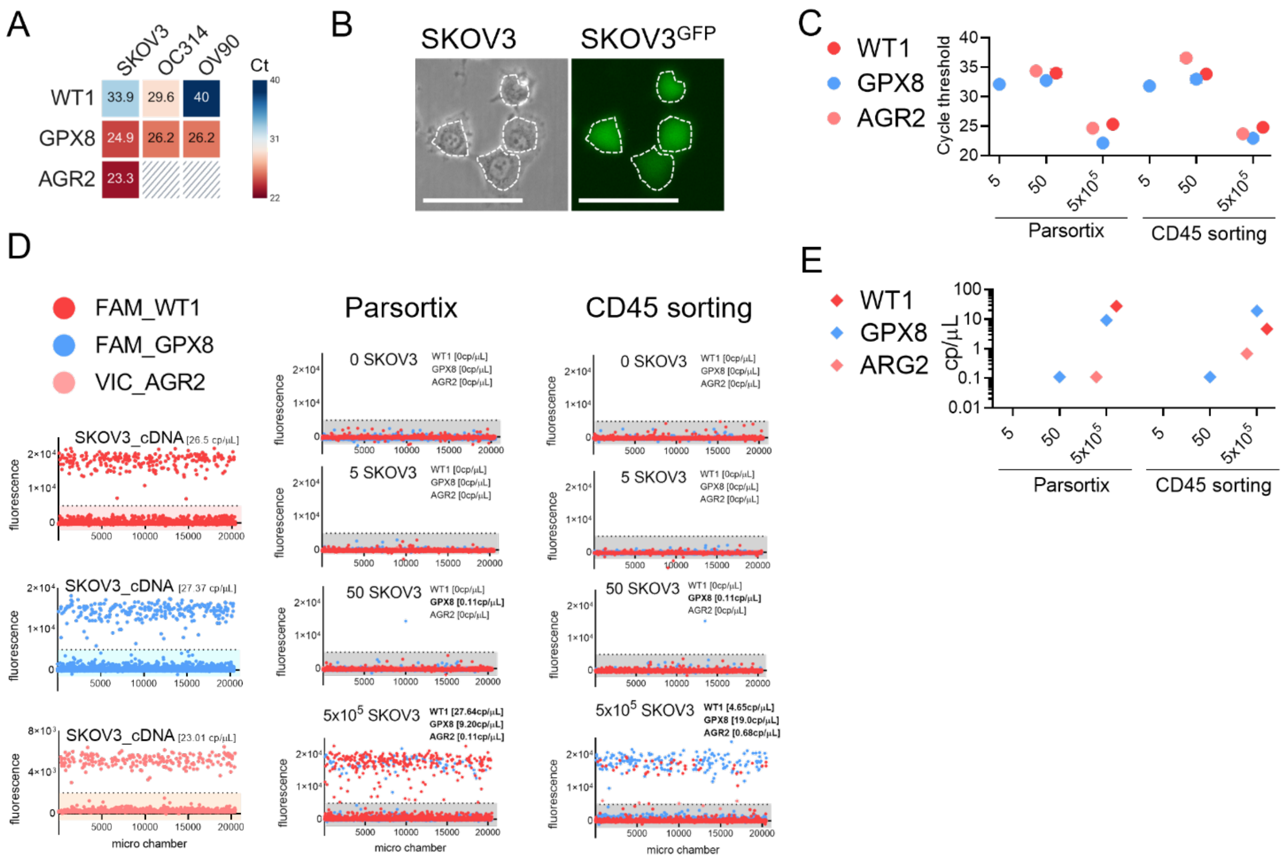


***Supplementary Figure 1. Analytical validity assessment of the ARG2/GPX8/WT1 three-gene assay to detect low cancer cell numbers in PB. (A)*** *Heatmap of Cycle thresholds (Ct) obtained via qRT-PCR of RNA extracted from three OC cell lines. Slashed boxes indicate undetectable transcripts.* ***(B)*** *Representative images of SKOV3 cells stably expressing GFP, acquired in phase contrast (left) and GFP fluorescence channel (right). Dashed lines highlight individual cells. Scale bar = 75 µm.* ***(C)*** *Dot plot reporting Cts obtained via qRT-PCR of preamplified (PA) cDNA derived from healthy donor PB samples spiked with 5, 50 and 500'000 SKOV3 cells and enriched with Parsortix or CD45 negative sorting.* ***(D)*** *Raw data extrapolated from QuantStudio Absolute Q dPCR software, indicating fluorescence signal on Y axis and microchamber location on X axis for WT1, GPX8 and AGR2. The dotted line indicates the detection threshold, while the computed cDNA concentration for each gene is indicated in brackets.* ***(E)*** *Dot plot reporting cDNA concentration obtained via dPCR of native cDNA derived from healthy donor PB spiked with 5, 50 and 500'000 SKOV3 cells enriched with Parsortix or CD45 negative sorting.*

A

| HGSOC- | *TP53* variant | ctDNA VAF (%) |
|---|---|---|
| 1 | c.376-2A>G | 1.38 |
| 2 | p.T253_I254del | 26 |
| 3 | p.Q167Hfs*3 | 0.5 |
| | p.Y205C | 0.4 |
| | p.Q52Mfs*4 | 0.21 |
| 6 | p.E294* | 0.2 |
| | p.P36S | 0.23 |
| 7 | p.G245D | 3.78 |
| 8 | p.L201Ffs*8 | 3.05 |
| 9 | p.G226Afs*r21 | 0.24 |
| 16 | p.G112Ffs*33 | 15 |
| 20 | p.R273H | 15.98 |
| | p.V272A | 0.27 |
| | p.K292E | 2.47 |
| | p.P278R | 0.23 |
| | p.Y220C | 0.27 |
| | p.H178P | 0.21 |
| 22 | p.Y234C | 1.03 |
| | p.R273H | 4.62 |
| | p.P300A | 1.77 |
| | p.K292E | 0.62 |
| 23 | p.R213Hfs*34 | 6.16 |
| 25 | p.C277F | 21.73 |
| 26 | p.P153Rfs*24 | 0.59 |
| | p.D184H | 0.28 |
| | p.R335H | 0.22 |
| 27 | p.W91* | 0.34 |
| 28 | p.G244C | 1.25 |
| 29 | c.560-1G>A | 0.84 |
| 30 | p.R248W | 3.06 |
| | p.N131Tfs*39 | 0.22 |

B

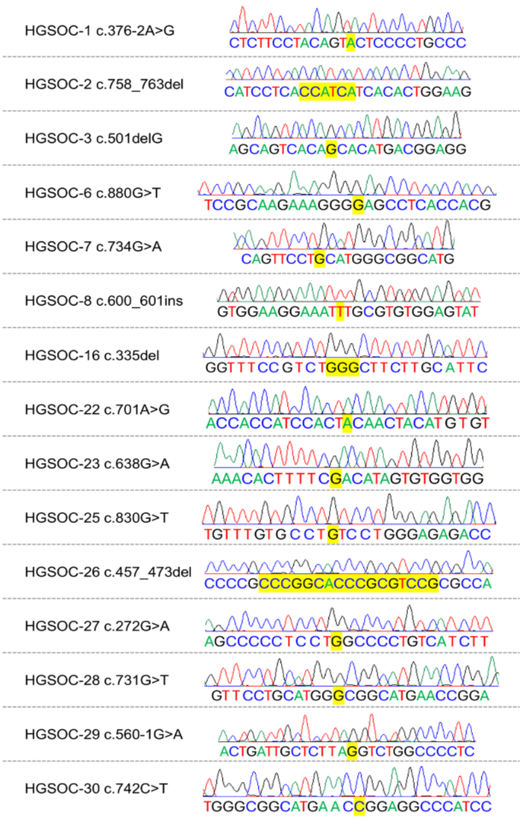


***Supplementary Figure 2. TP53 sequencing results used for setting the Variant Allele Frequency threshold for ctDNA mutational call. (A)*** *Table listing all TP53 variants found in cfDNA by high-depth TP53 sequencing prior optimal VAF threshold application. Mutations found in primary tumor biopsy are indicated in black, whereas variants undetected in primary tumor are indicated in red.* ***(B)*** *PBMC sequencing results confirming somatic nature of primary tumor mutations. For each HGSOC patient the TP53 mutation is reported, together with corresponding Sanger sequencing chromatograms of matching PBMC, where wild-type genotype is indicated in yellow. GRCh37 - ENST00000269305.4 TP53-001 was used as reference.*
